# PyXlinkViewer: a flexible tool for visualisation of protein chemical crosslinking data within the PyMOL molecular graphics system

**DOI:** 10.1101/2020.06.16.154773

**Authors:** Bob Schiffrin, Sheena. E. Radford, David. J. Brockwell, Antonio N. Calabrese

## Abstract

Chemical crosslinking-mass spectrometry (XL-MS) is a valuable technique for gaining insights into protein structure and the organization of macromolecular complexes. XL-MS data yields inter-residue restraints that can be compared with high-resolution structural data. Distances greater than the crosslinker spacer-arm can reveal lowly-populated “excited” states of proteins/protein assemblies, or crosslinks can be used as restraints to generate structural models in the absence of structural data. Despite increasing uptake of XL-MS, there are few tools to enable rapid and facile mapping of XL-MS data onto high-resolution structures or structural models. PyXlinkViewer is a user-friendly plugin for PyMOL v2 that maps intra-protein, inter-protein and dead-end crosslinks onto protein structures/models and automates the calculation of inter-residue distances for the detected crosslinks. This enables rapid visualisation of XL-MS data, assessment of whether a set of detected crosslinks is congruent with structural data, and easy production of high-quality images for publication.

## 1. Introduction

Chemical crosslinking-mass spectrometry (XL-MS) is a powerful tool to derive structural information on proteins/protein assemblies, including transient or weak interactions necessary for biological function (1–3). XL-MS workflows begin by treatment of a purified protein/protein complex, lysate or intact cell with an appropriate XL reagent. All crosslinkers have the same basic architecture, comprising two reactive groups (e.g. NHS-esters that primarily react with amines (4)), or radical based crosslinkers (such as diazirines (5) that react non-specifically) separated by a spacer arm (1–3). This spacer arm confers a distance constraint that can be used for comparison with high-resolution structures, structural modelling, model validation or the detection of lowly-populated states not captured by other methods.

After crosslinking, the proteins are digested (e.g. with trypsin) and the inter-protein, intra-protein and dead-end (where only one of the two reactive groups react with the protein and the other remains unreacted or is quenched by solvent) crosslinked peptides must then be detected. These peptides are of relatively low abundance compared to their non-crosslinked counterparts, so much work has focused on developing analytical workflows to enable crosslink detection (e.g. using enrichment protocols (6), incorporating MS-cleavable groups in the spacer arm that yield diagnostic fragment ions (7), and developing specialised informatics tools (8)). The detected crosslinks can be mapped onto protein sequences/networks (e.g. using xVis (9) or xiNET (10)) or protein structures for comparison with other data. Current tools for XL-MS data visualisation include XLinkAnalyzer (11), a plugin for Chimera (12), as well as tools for the calculation and visualisation of crosslinks as solvent accessible surface distance (SASD) paths (Xwalk (13), Jwalk (14) and DynamXL (15)). However, no plugin exists for the commonly used PyMOL molecular graphics system.

Here we present PyXlinkViewer, a tool for mapping chemical crosslinks onto a protein structure or structural model visualised within the PyMOL v2 molecular graphics system. PyXlinkViewer automates the display of inter-protein and intra-protein crosslinks and measures the Cα-Cα inter-residue Euclidean distances for each crosslink. These values can then be compared with the upper distance limit imposed by the crosslinker used. PyXlinkViewer also displays residues modified with dead-end crosslinks, which can probe for chemical accessibility and be used to score structural models (16), hence our tool can also be used to easily visualise data from a range of other covalent labelling/footprinting workflows (e.g. fast photochemical oxidation of proteins (17), carbene labelling (18) or other side-chain-specific probes (19)). We envisage that this versatile tool will be a useful addition to XL-MS and other covalent labelling-MS workflows.

## 2. Results

We have developed the PyXlinkViewer plugin to enable rapid, simple visualisation of inter-residue chemical crosslinks and dead-end crosslinks mapped onto a protein structure or structural model visualised within PyMOL v2. The source code, plugin installation file, example data, and user manual are freely available under a GNU General Public License at https://github.com/BobSchiffrin/PyXlinkViewer.

### 2.1 Plugin design and installation

PyXlinkViewer is a cross-platform plugin for PyMOL v2 written in Python 3, which can be run on Linux, macOS, or Microsoft Windows operating systems. It has a dedicated graphical user-interface (GUI) (**Figure 1**) designed using QtDesigner v4.8.6. PyXlinkViewer is supplied as a ZIP file, with installation easily achieved using PyMOL’s plugin manager. A detailed User Manual is available with the software. The code only imports modules present in the standard PyMOL v2 and Python 3 libraries, so no external dependencies need to be installed.

**Figure 1.**
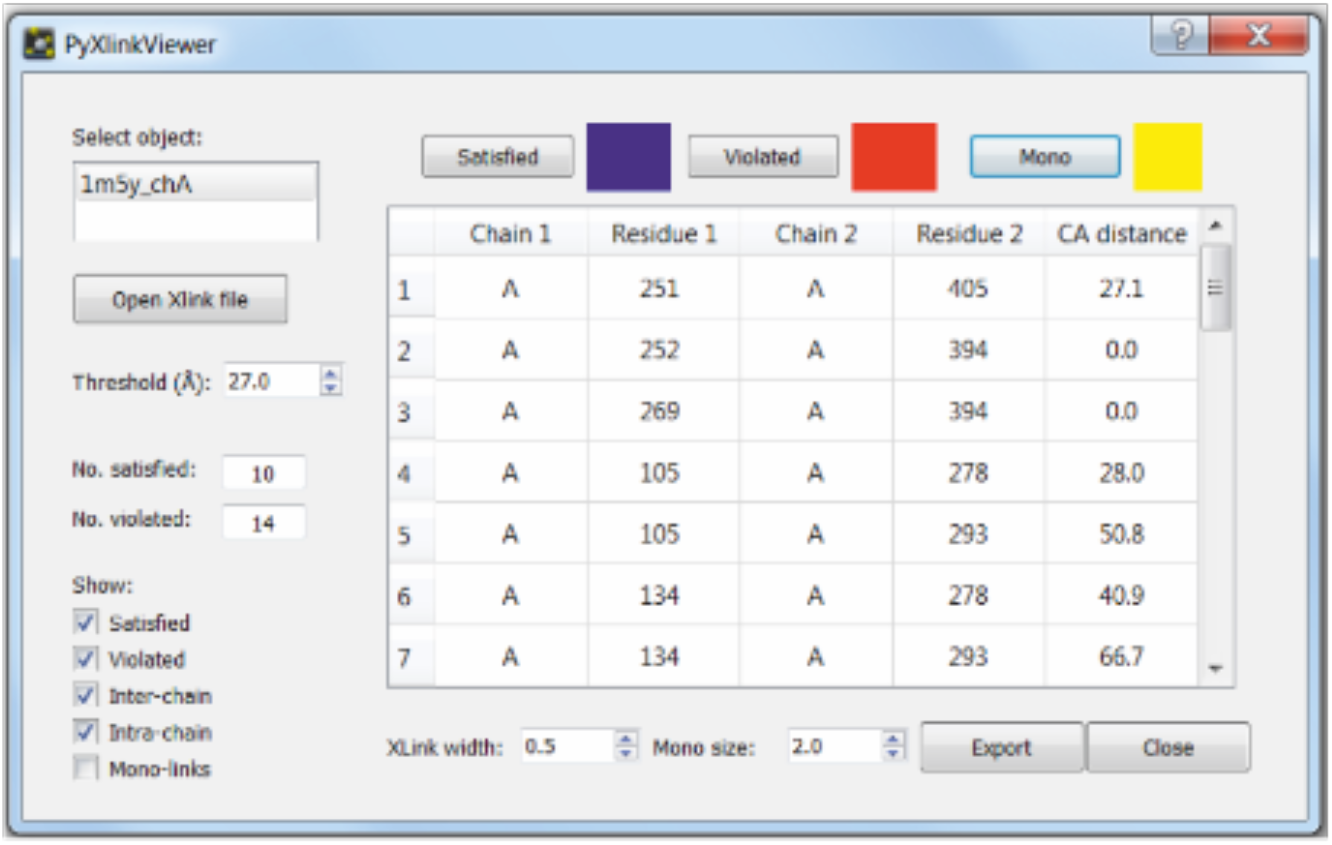
PyXlinkViewer graphical user interface. A PyMOL object containing the protein structure(s) for which the crosslinking data is to be visualised is first selected (top left). The crosslinking data file is then opened and the crosslinks are displayed on the structure in the PyMOL viewer and in the table in the PyXlinkViewer GUI. The user can edit the threshold value used to determine if a crosslink is satisfied or violated (determined from the structure of the crosslinking reagent). Check boxes control the display of various crosslink types (satisfied, violated, inter-chain, intra-chain, mono-links) in both the PyMOL viewer and in the table in the PyXlinkViewer GUI. The colours and sizes of the displayed crosslinks can be easily modified and a data export function is included.

### 2.2 Data import

After manual curation of an appropriate XL-MS dataset using dedicated XL-MS analysis software (8), the data are prepared as a list of crosslinks and dead-ends for use in PyXlinkViewer. This approach enables the user to curate their dataset and filter it based on the score of the detected crosslinks (each data analysis software package uses a different scoring algorithm) prior to data visualisation. The file format required is that used by the existing tool Jwalk (14) (see User Manual) and allows maximum user control of the data displayed (see example data). This format details the residue numbers and chain identifiers of both residues involved in a crosslink. To visualise dead-end or covalent labelling/footprinting data, only one residue/chain identifier is included in each entry (see example data). There are a number of different file formats generated by the various XL-MS data processing tools (8). However, recently a XL-MS file conversion utility has been created (20), and a standard XL-MS data format proposed. The object-orientated design of PyXlinkViewer allows for simple extension to support a standard file format if adopted by the XL-MS community.

### 2.3 Data visualisation using PyXlinkViewer

After opening a data file, a table of crosslinks is populated and the inter-residue Cα-Cα Euclidian distances are calculated and displayed (**Figure 1**). The upper distance threshold imposed by the spacer arm can be set, and the number of crosslinks that are satisfied/violated by this distance threshold is displayed. Crosslinks and dead-ends are drawn as lines between residue Cα atoms, or spheres at residue Cα atoms, respectively. Often residues are missing from the PDB files of protein structures (e.g. if they are disordered/unresolved by X-ray crystallography or cryogenic electron microscopy). If a residue involved in a crosslink is not present in the structure loaded into PyXlinkViewer, a warning message is printed in the PyMOL command window that the crosslink cannot be shown, which may draw attention to disordered regions involved in a protein interaction.

A number of visualisation options can be changed in the GUI, including the colour and width of the drawn crosslinks. The user can also show or hide different crosslink types (i.e. inter-protein, intra-protein and dead-end). The XL table in the PyXlinkViewer GUI and the data visualised in the PyMOL display are automatically updated when the user makes any changes to the visualisation options (see User Manual). Each XL is drawn as a separate PyMOL object, labelled with the chain and residue IDs involved, so can be individually shown/hidden using the PyMOL GUI or command line. The data can be exported as a CSV file for further interrogation.

### 2.4 Visualising XL-MS data for monomeric proteins and multimeric assemblies

To demonstrate the power of PyXlinkViewer, we highlight here its use with two publically available XL-MS datasets.

Firstly, we demonstrate the PyXlinkViewer workflow using intra-protein crosslinks detected in the chaperone SurA (**Figure 2**). SurA comprises three domains, a core domain (made of an N-terminal and C-terminal region), and two PPIase domains, P1 and P2 (**Figure 2**). SurA was crosslinked with the reagent DSBU, which contains two NHS-ester reactive groups (that primarily react with Lys residues and protein N-termini, but also with Ser, Thr, Tyr), and has been shown to crosslink residues up to ~27-30 Å apart (Cα-Cα Euclidean distances) (21). Of the 32 intra-protein crosslinks detected, 13 were incompatible with the crystal structure when using a Euclidian distance cut-off of 27 Å, as determined using PyXlinkViewer. 8 crosslinks in SurA involved residues missing from the PDB structure (as detected by PyXlinkViewer) which were subsequently built using MODELLER (22). As a PyMOL object is created for each crosslink, intra-domain and/or inter-domain crosslinks could easily be individually selected for display. While only 2/13 intra-domain crosslinks were incompatible with the crystal structure, most of the inter-domain crosslinks (11/19) were between residues greater than 27 Å apart. The data provided evidence that in solution SurA populates conformations in which the P2 domain is much to the core and P1 domains than suggested by the crystal structure (23; 24).

**Figure 2.**
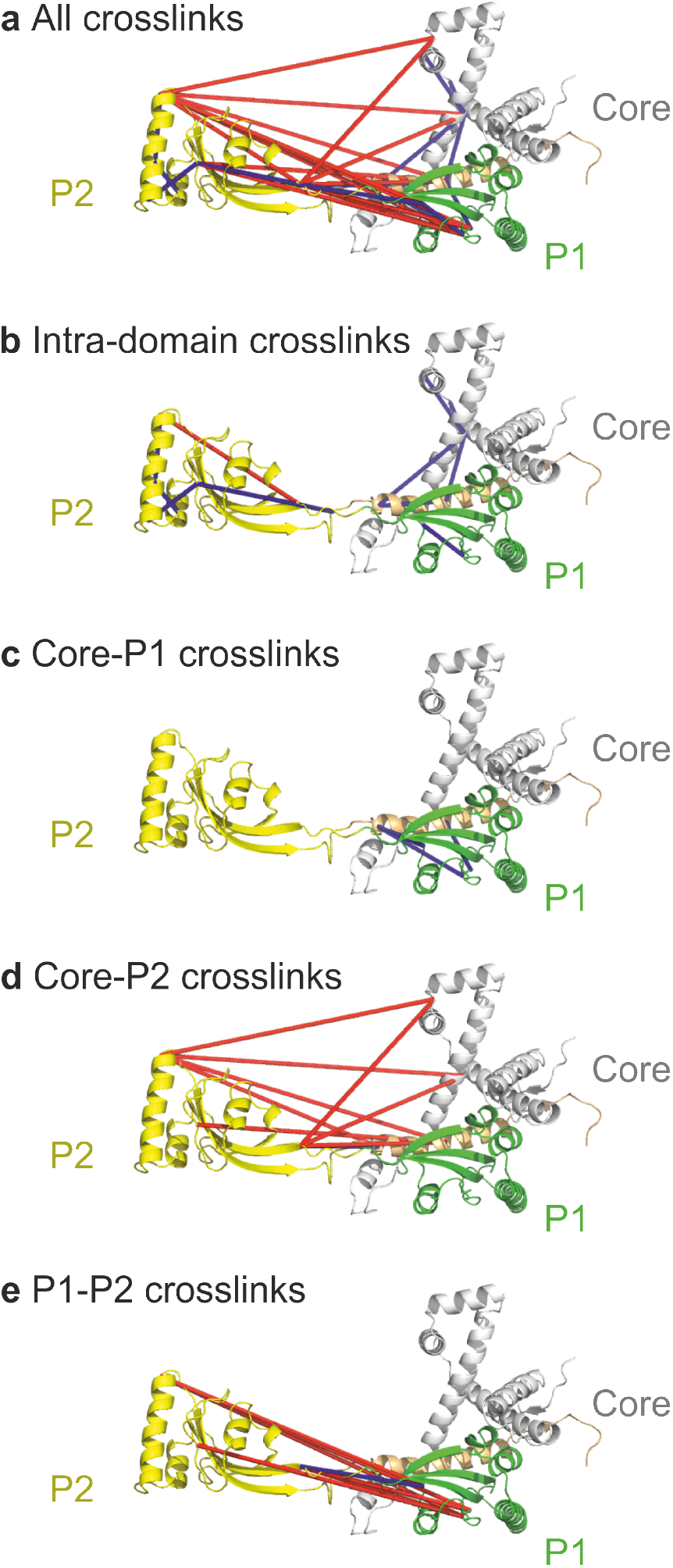
Visualisation of SurA intra-protein crosslinks using PyXlinkViewer. SurA has a three domain architecture, comprising the core domain (grey/orange), and two PPIase domains, P1 (green) and P2 (yellow). In the crystal structure of SurA (PDB: 1M5Y), the P2 domain is spatially separated from the core and P1 domains (23). **(a)** All detected crosslinks. **(b)** Intra-domain crosslinks. **(c-e)** Crosslinks detected between the **(c)** Core-P1, **(d)** Core-P2, and **(e)** P1-P2 domains. These data suggest that the P2 domain populates conformations in solution closer to the core and P1 domains than suggested by the crystal structure. Crosslinks are coloured blue or red, if they are satisfied or violated, respectively, using a Cα-Cα Euclidean distance threshold of 27 Å.

In order to demonstrate PyXlinkViewer’s functions on a larger, multicomponent complex, we chose to visualise a XL-MS dataset obtained for the OCCM complex (**Figure 3a**), a helicase loading intermediate in DNA replication (14 subunits, ~1 MDa, 1132 Lys-Lys crosslinks) (25; 26). On mapping the crosslinks to the cryoEM structure of OCCM (25), PyXlinkViewer identified that many of the crosslinks (625/1132) involved residues that were not present in the cryoEM structure. Of those that could be displayed, 77/507 were between residues greater than 27 Å apart in the structure. Using PyXlinkViewer’s ability to selectively display intra-chain or inter-chain XLs (**Figure 3b,c**), it was easily possible to see that the violated intra-chain XLs are between the domains of multi-domain protein components of OCCM. Notably, violated intra-protein crosslinks were detected in the three proteins Cdc6, Mcm3 and Cdt1, suggesting some inter-domain motions/reorganisation in these proteins in the complex (**Figure 3c**) Additionally, it was easily discernible that many of the violated inter-protein XLs involved regions of different proteins within the complex that are not in defined secondary structure. Combined, these two examples of SurA and OCCM demonstrate that PyXlinkViewer can be used to visualise both large and small XL-MS datasets to rapidly gain structural/functional insights into proteins and their assemblies.

**Figure 3.**
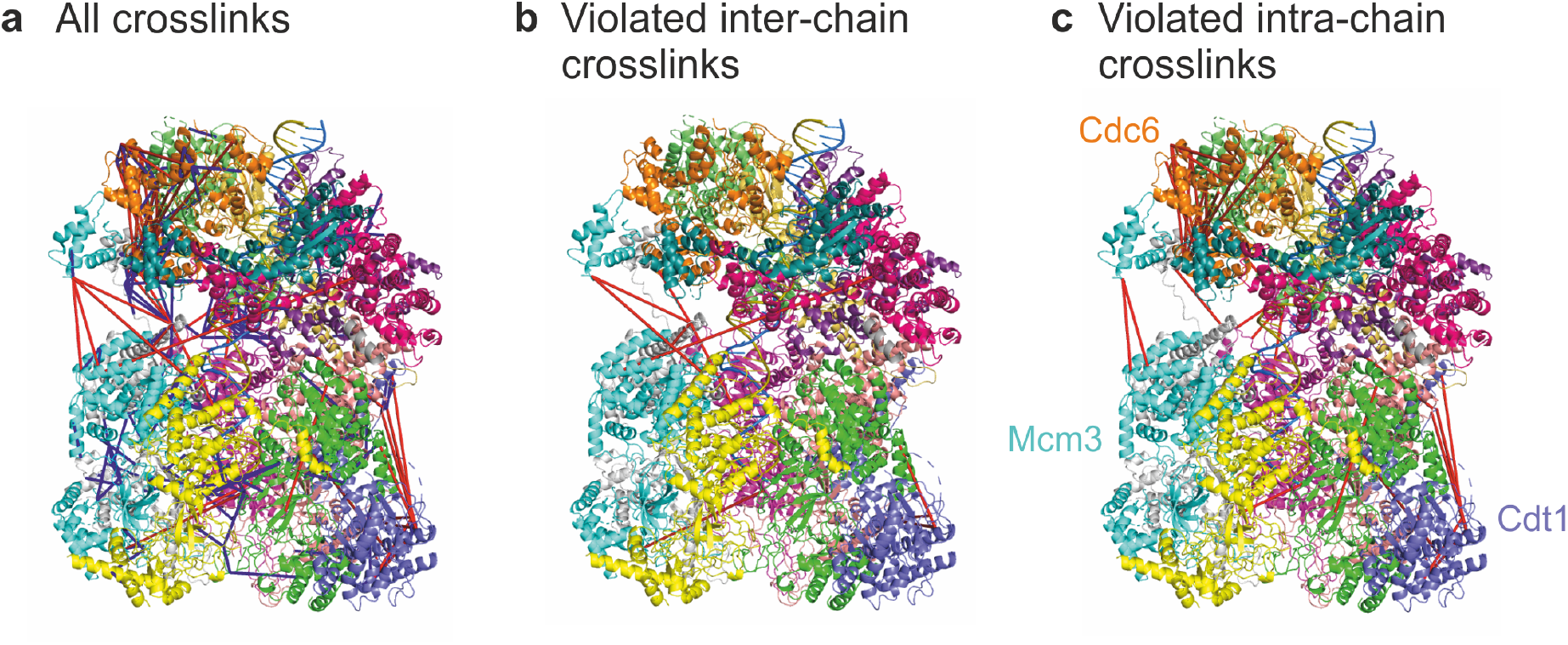
PyXlinkViewer enables rapid visualisation of 507 crosslinks within the 14 component ~1 MDa OCCM complex (PDB: 5V8F (25)). 1132 crosslinks were detected in the dataset (26), 625 of which involved residues that were not present in the PDB structure and are therefore not shown here. **(a)** All crosslinks mapped to the structure of OCCM. **(b)** Violated inter-chain crosslinks. (**c)** Violated intra-chain crosslinks. Crosslinks are coloured blue or red, if they are satisfied or violated, respectively, using a Cα-Cα Euclidean distance threshold of 27 Å. Each subunit is shown in a different colour in this figure. Three subunits are labelled in (c) as the intra-protein crosslinks detected here suggest some inter-domain motions.

## 3. Discussion

Recent enhancements in XL-MS protocols (6; 27; 26), new crosslinker designs (28), and refined bioinformatics approaches (8), have ensured XL-MS is a key tool in the structural biologist’s armory of methods for determination of the structure and dynamics of proteins and their complexes both *in vitro* and *in vivo*. Computational modelling methods using XL-MS restraints are being constantly improved (15; 29; 14; 16), but a vital, final step in this pipeline is the comparison of structures/structural models with XL-MS data. This is necessary for model validation and data presentation. PyXlinkViewer is the first PyMOL plugin specifically designed to simply map crosslinks onto high-resolution structures/structural models, which is key for increasing the throughput of integrative structural studies. For reasons of speed and clarity, PyXlinkViewer calculates and displays Euclidean distances for each inter-residue XL. However, solvent accessible surface distances (SASDs) can be a more reliable indicator of whether or not a cross-link is satisfied (13; 14), and routines which take into account side-chain dynamics have also been developed (15). However, calculation of SASDs is currently ~5 orders of magnitude slower than for Euclidian distances (14), potentially prohibiting this approach when dealing with large numbers of crosslinks and/or structural models. The object-orientated design of PyXlinkViewer allows for extension of functionality in the future, including possible calculation and display of SASDs.

Discriminating between inter- and intra-subunit crosslinks in homooligomeric complexes remains an area of significant challenge. PyXlinkViewer allows the user to explicitly specify PDB chain identifiers for each residue pair in the input file, therefore all possible crosslinks can be displayed on the structure. This gives the user an opportunity to visually compare the distances of possible inter- and intra-subunit crosslinks for the same residue pair from different subunits and assess which are most compatible with the crosslinked peptides detected by MS. Experiments involving the detection of crosslinks between ‘light’ (^14^N) and ‘heavy’ (^15^N-labeled) subunits in a homooligomer, can also assist in discriminating between inter- and intra-subunit crosslinks (30–32).

PyXlinkViewer also enables visualisation of dead-end crosslinks and data from other covalent labelling workflows (e.g. those that target specific amino acid residues (19), or radical labeling methods such as fast photochemical oxidation of proteins [FPOP] which labels promiscuously (33; 17)). A number of software tools have been designed to enable the identification of residues that have been covalently labeled (34; 35). The data from these analyses can easily be converted to a format compatible with import to PyXlinkViewer. However, it should be noted that for reagents/techniques that label many types of residues, e.g. FPOP, it remains analytically challenging to determine at the residue level the site of the modification. Nevertheless, data from covalent labelling has shown promise as an input for structural modelling (36–38). To demonstrate the data input requirements for covalent labelling/ dead-end XL data we have also included example data from FPOP analysis of the protein β_2_-microglobulin (17) (see example data). Whilst dead-end crosslinking data are often discarded from analysis, especially when XL-MS is used to aid structural modelling, there is emerging evidence of their importance as probes for chemical accessibility. Indeed they can be used as restraints in modelling pipelines and as a tool for the scoring of structural models of proteins (36; 16).

In conclusion, PyXlinkViewer is a versatile plugin for PyMOL v2 that enables rapid interrogation of a XL-MS dataset by mapping crosslinks onto a structure and calculating their inter-residue distances. As PyXlinkViewer functions within PyMOL v2, publication quality figures depicting XL-MS data can be made with ease. We envisage that PyXlinkViewer will be a key tool for the expanding number of researchers using XL-MS to interrogate the functional mechanisms of proteins and protein assemblies.

## Acknowledgements

Members of the Radford and Brockwell labs are thanked for helpful discussions. This work has been supported by the BBSRC (BB/T000635/1, BB/P000037/1, BB/N007603/1). ANC acknowledges support from a University Academic Fellowship from the University of Leeds.

## Availability

The source code, plugin installation file, test data and user manual are freely available under a GNU General Public License at https://github.com/BobSchiffrin/PyXlinkViewer. PyMOLv2 is available from https://pymol.org/2/.

